# GC-1 spg cells are most similar to Leydig cells, a testis somatic cell-type, and not germ cells

**DOI:** 10.1101/2021.01.07.425754

**Authors:** Andrew R. Norman, Lauren Byrnes, Jeremy F. Reiter

## Abstract

GC-1 spg is an immortalized cell line derived from an adult mouse testis and reported to be most similar to spermatocytes, a male germ cell-type. However, immunofluorescence indicates that GC-1 spg cells express WT1, a marker of testis somatic cells, and do not express markers of germ cells. Transcriptomic profiling indicate GC-1 cells are most similar to Leydig cells. Therefore, we conclude that GC-1 spg cells are most similar to testis somatic cells.

## Introduction

The testis is a complex organ consisting of tissues derived from multiple lineages. At day 11.5 in the mouse, pre-Sertoli cells migrate from the coelomic epithelium to the developing gonad and aggregate with primordial germ cells. Together with surrounding interstitial and endothelial cells, these aggregates form the sex chords, which mature into seminiferous tubules, the sperm-producing structures of the testis. Further differentiation of the interstitial cells produces Leydig cells, the principle source of testosterone in males (*1*).

This complexity has made it difficult to entirely recapitulate testes development in vitro for the purposes of study or sperm production, although partial success has been achieved with germ-cell co-culture systems (*2*) and testes ex-vivo culture (*3, 4*). To study a single testes cell type, immortalized testes cell lines can be tractable. To this end, Hofmann et al. derived several SV40 T antigen-immortalized cell lines from adult mouse testis (*5*). Based on morphology and expression of a spermatocyte-specific marker, one of these cell lines, called GC-1 spg or GC-1, was described as being similar to spermatocytes, a male meiotic germ cell-type (*6*). Using transcriptome analysis and immunocytochemistry, we conclude that GC-1 cells are that GC-1 cells are more to Leydig cells, a testis somatic cell-type, than to any germ cell-type.

## Results

If GC-1 cells are similar to spermatocytes, a germline cell-type, then these cells should express germline-specific proteins. DDX4 is a germline-specific RNA helicase, and an established marker of germline cells in both Drosophila and mammals (*7*). Consistent with prior results, immunofluorescence revealed that the luminal germ cells of mouse testis tubules expressed DDX4. In contrast, GC-1 cells did not express DDX4, similar to the fibroblast NIH/3T3 cells (Fig. 1,A-C).

**Figure 1.**
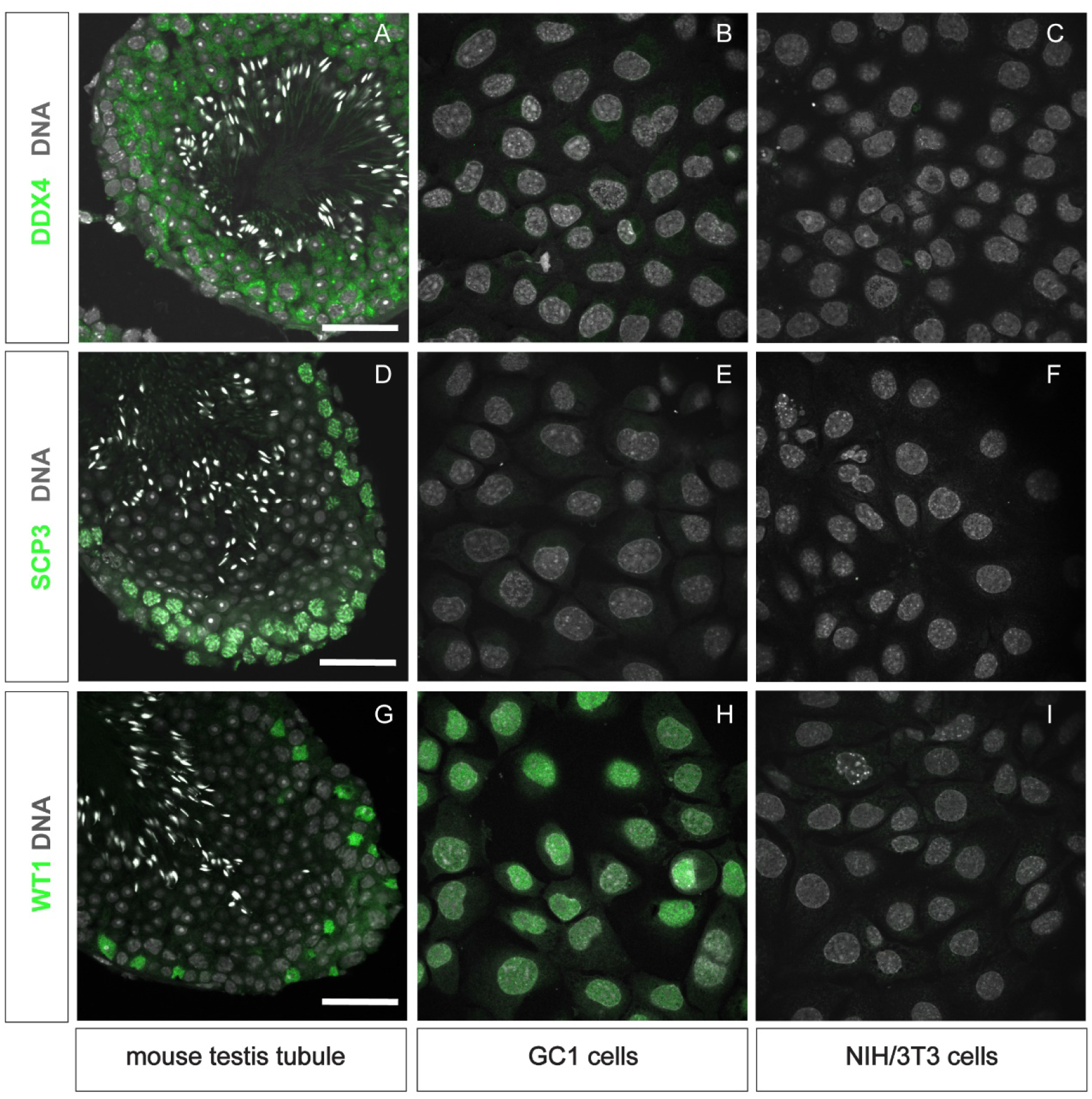
GC-1 cells express a marker of testis somatic cells, and not markers of germline cells. (A-C) DDX4 (green) staining of mouse testes, GC-1 cells and NIH/3T3 cells. DDX4 is not expressed by GC-1 or NIH/3T3 cells, but is expressed by the luminal testis germline cells in the germinal epithelium. (D-F) SCP3 (green) staining. SCP3 is not expressed by GC-1 or NIH/3T3 cells. (G-I) WT1 (green) staining. Sertoli cells in the testis (G) and GC-1 cells (H) express WT1, but not NIH/3T3 cells (I). Hoechst (gray) marks nuclear DNA. Scale bars are 50 µm.

SCP3 is a component of the meiotic synaptonemal complex expressed by spermatocytes (*8*). Spermatocytes of the mouse testis, recognized by their large nuclei and proximity to condensing post-meiotic spermatids, expressed SCP3 (Fig. 1,D-F). Again, in contrast, GC-1 cells did not express SCP3, similar to NIH/3T3 cells.

WT1 is a transcription factor expressed by Sertoli cells and some testis interstitial cell precursors, including those of Leydig cells (*9*) (Fig. 1G). GC-1 cells did express WT1, whereas NIH/3T3 cells did not (Fig. 1 H-I). Taken together, these immunohistochemical findings suggest that GC-1 cells are more similar to testis somatic cells than to germ cells.

To confirm the identity of the GC-1 cells, we compared the GC-1 transcriptome with the transcriptomes of known mouse testis cell-types, as reported in a recent study using single-cell RNA sequencing (*10*). We utilized the data analysis pipeline and cell clustering identification metadata of Green et al. to reconstruct the whole-testis dataset (Fig. 2A). We then determined protein-coding transcript read counts of GC-1 cells by bulk RNA-sequencing.

**Figure 2.**
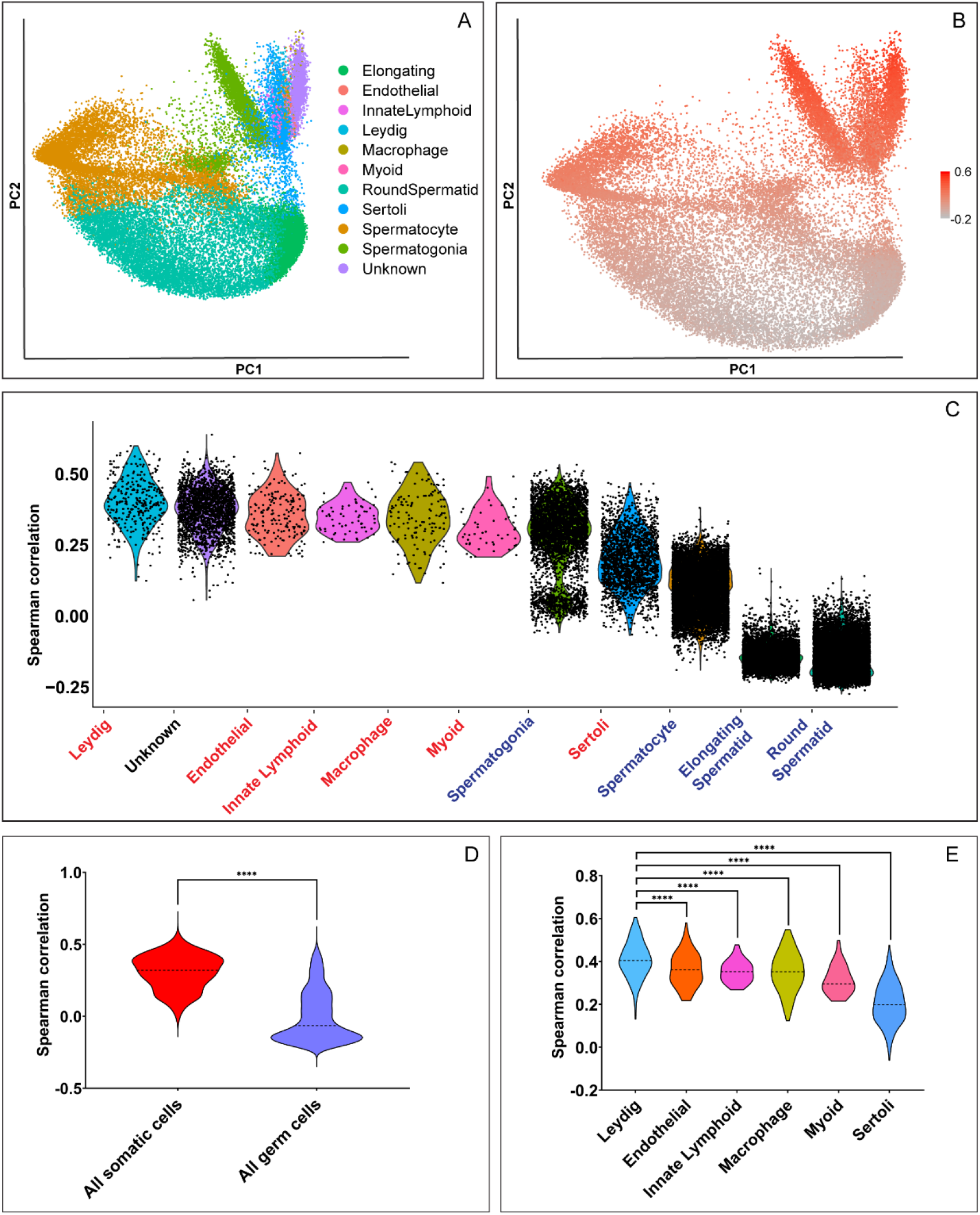
GC-1 cells are transcriptionally more similar to testis somatic cells than to testis germ cells. (A) Reconstruction of principle components of data from Green and colleagues *(5)* (B) Spearman correlation between the transcriptomes of the cell types reported by Green and colleagues and the transcriptome of GC-1 cells determined by bulk sequencing. Color intensity is proportional to correlation of transcriptomes. (C) The Spearman correlations between the GC-1 transcriptome and each testis cell type designated in the single-cell dataset. Black dots signify individual cell transcriptomes.The same data are presented but re-grouped by germ cell or somatic testis cell identity(D) and by specific somatic cell identity (E). Dotted lines in violin plots mark population mean. P-values from Student’s t-test.

Sequencing of single-cell RNAs is more shallow and sparse than that of bulk RNA, complicating direct comparison of the two kinds of data. We reasoned that the rank order of gene expression in the two kinds of data should be unaffected by depth of sequencing, especially for more highly expressed genes. Therefore, we used a Spearman correlation to compare the rank order of gene expression of the bulk GC-1 transcriptome to the rank order of gene expression for each of the testis single-cell transcriptomes (Fig. 2 B-C). To ensure comparison of highly expressed genes from the single-cell dataset, we used the cluster marker genes identified by Green et al. for the correlation analysis. Multiple somatic cell types correlated more highly with the GC-1 transcriptome than the germline cell types (Fig. 2C-D). In particular, the GC-1 cells were found to be more similar to Leydig cells than to any identifiable testis cell-type (Fig. 2E).

## Discussion

GC-1 cells were previously reported to be most similar to spermatocytes (*5*). For example, GC-1 cells were reported to express lactate dehydrogenase C4 (LDHC) and testis-specific cytochrome C (CYCT) based on immunocytochemistry (*5*). Here, through complementary immunocytochemical and transcriptomic analyses, we have shown that GC-1 cells are more similar to testis somatic cells than to germ cells. For example, transcriptomic analysis reveals that GC-1 cells do not express *Ldhc* or *Cyct*, suggesting that prior immunocytochemical staining was non-specific. While the transcriptomic data indicate that GC-1 cells are most similar to Leydig cells, WT1 expression is not typically a feature of adult Leydig cells. However, WT1 is expressed in the shared precursors of Leydig and Sertoli cells (*9, 11*). It is most likely that GC-1 cells, a highly proliferative cell line, are derived from a proliferating Leydig cell precursor population.

## Methods

### Cell Culture

Mouse GC-1 testes cells and NIH 3T3 cells were cultured in Dulbecco’s Modified Eagle’s Medium (Glutamax and HEPES supplementation; Invitrogen Cat# 10564) supplemented with Pen/Strep (10,000 U/mL; Invitrogen Cat# 15140148). Medium was changed every 2 days and cells were passaged upon 90% confluency using Trypsin-EDTA (0.05%; Invitrogen Cat# 25300054). Cells were maintained at 37°C in a 5% CO2 atmosphere.

### Immunofluorescence

GC1 and 3T3 cells were grown to confluency on glass cover slips and fixed in 4% paraformaldehyde for 10 minutes. Adult mouse testes were fixed in 4% paraformaldehyde overnight at 4°C, equilibrated in PBS with 30% sucrose overnight at 4°C, embedded in O.C.T. Compound (Tissue-Tek) and sectioned with a cryostat (model?) at 12 microns. Cells and mouse testes sections were blocked in PBS with 0.1% Triton X-100/2% BSA/1% normal donkey serum for 1 hour at 4°C and stained with primary antibodies (see below) overnight at 4°C in the blocking solution. Cells and mouse testes sections were stained with secondary antibodies and Hoechst 33342 (1µg/mL) in PBS with 5% normal donkey serum for 1 hour at room temperature. Processing of cells and mouse testes sections was done identically and in parallel. For staining with the DDX4 and SCP3 antibodies, an antigen retrieval was performed by incubating the fixed slides in a sodium citrate buffer (10mM sodium citrate and 0.05% Tween-20, pH 6.0) for 20 minutes at 95°C, prior to the antibody staining protocol. Coverslips were mounted on glass slides in Prolong Diamond mounting medium (Life Technologies) and imaged on a Leica TCS SPE DM5500Q confocal microscope. An ACS Apochromat 63X objective (1.30 NA, oil immersion) was used with the standard Leica LAS AF acquisition software.

### Antibodies

Mouse anti-SCP3 (Abcam, ab97672, 1/250), rabbit anti-WT1 (Abcam, ab89901, 1/100) and rabbit anti-DDX4 (Proteintech, 51042-1-AP, 1/500) were used as primary stains. Donkey anti-Mouse/Rabbit IgG (H+L) Highly Cross-Adsorbed Secondary Antibody conjugated to Alexa 488 (Life Technologies, 1/2000) were used to detect the primary antibody staining.

### Bulk RNA sequencing

GC1 cells were grown to 90% confluency in triplicate, then lysed directly in buffer RLT according to the standard RNeasy kit protocol (Qiagen). Lysate was homogenized with the QIAshredder columns (Qiagen) and processed according to the standard RNeasy kit protocol. RNA integrity was confirmed with a BioAnalyzer (Agilent) and a sequencing library was constructed using the TruSeq PolyA kit (Illumina). Sequencing was performed with an Illumina HiSeq at a length of 50bp. Read alignment was performed using STAR (v2.6.0), with the mouse GRCm38.88 genome and default settings (Dobin et al., 2013). Read counts per coding gene (TPM values) were determined using RSEM (v1.2.25) with the default settings (Li et al., 2011).

### Single-cell Analysis and Bulk Dataset Correlation

Scripts from https://github.com/qianqianshao/Drop-seq_ST and data from GSE112393 were downloaded and processes as described in Green et al.. The genes used for correlation were those determined by Green et al. to be markers of the 11 cell types identified in the single-cell dataset (Table S2). The log(UMI count + 1) for each single cell in the single-cell dataset was then correlated to the log(TPM + 1) of the bulk dataset using Spearman rank correlation. Scripts for correlation analysis and plotting are available at https://github.com/lb15/GC1_paper.

